# Phigaro: high throughput prophage sequence annotation

**DOI:** 10.1101/598243

**Authors:** Elizaveta V. Starikova, Polina O. Tikhonova, Nikita A. Prianichnikov, Chris M. Rands, Evgeny M. Zdobnov, Vadim M. Govorun

**Affiliations:** Department of Molecular Biology and Genetics, Federal Research and Clinical Centre of Physical-Chemical Medicine, Malaya Pirogovskaya 1a, 119435 Moscow, Russia; Department of Genetic Medicine and Development, University of Geneva Medical School and Swiss Institute of Bioinformatics, Geneva, Switzerland

## Abstract

**Summary:** Phigaro is a standalone command-line application that is able to detect prophage regions taking raw genome and metagenome assemblies as an input. It also produces dynamic annotated “prophage genome maps” and marks possible transposon insertion spots inside prophages. It provides putative taxonomic annotations that can distinguish tailed from non-tailed phages. It is applicable for mining prophage regions from large metagenomic datasets.

**Availability:** Source code for Phigaro is freely available for download at https://github.com/bobeobibo/phigaro along with test data. The code is written in Python.

## Introduction

Bacteriophages (phages) are viruses that infect bacteria and have recently gained increasing interest due to the alarming spread of antibiotic-resistant strains of pathogenic bacteria. Phages are known for their substantial impact on diverse ecosystems, from animals’ intestinal tracts to oceans. Phages can sometimes provide benefits to their hosts by transporting virulence factors and antibiotics resistance genes among bacterial strains. To date, our knowledge of bacteriophage diversity is narrow due to a negligible number of isolated and sequenced bacteriophage genomes, as compared to the huge proportion of viral “dark matter” found in metagenomes [11]. Many undiscovered viral sequences of Myoviridae, Podoviridae, Siphoviridae, Inoviridae and Microviridae families lie within sequenced bacterial genomes in the form of prophages, as those families are known to have temperate life cycles, and even more unknown prophages are likely within metagenomes. Existing command line tools for prophage prediction tend to output a limited selection of annotations and visualizations, and generally don’t mark any overlapping mobile elements like transposons. Here we present Phigaro, a novel high-throughput command line tool that is able to predict and annotate prophage sequences with a dynamic visualization interface applicable to both genomic and metagenomic assembled data.

## Phigaro overview

Phigaro is a Python package that accepts one or more FASTA files of assembled contigs as input. The core of this program is PhigaroFinder algorithm that defines regions of putative prophages based on preprocessed input data. The preprocessing is carried out consistently by two external programs. Firstly, FASTA files are processed by Prodigal [6], which returns a list of genes with their coordinates, GC content and other properties for a given sequence. Then the obtained genes are annotated with HMMSCAN [9] using phage-specific profile HMMs from pVOGs (prokaryotic Virus Orthologous Groups) [5]. A gene is considered “phage-like” if it corresponds to one of the pVOG profile HMMs.

### PhigaroFinder algorithm

For each gene, PhigaroFinder algorithm computes the probability of it being localized in a prophage region. The algorithm uses two pre-computed sets of pVOG profile HMMs: the “black list” and the “white list”. Those lists were formed based on pVOG distributions inside and outside known prophage regions in 54 bacterial genomes in order to correct the initial set of pVOG profile HMMs to avoid detecting regions with a high density of genes corresponding to pVOGs that are, in fact, not true prophage regions. The “black list” consists of pVOGs that are likely to be found in other regions unrelated to prophages throughout bacterial genomes (e.g. the ones annotated as “ABC transporters”, “plasmid partition proteins”, etc.), while the “white list” is the opposite: it consists of pVOGs that are more likely to be found in prophage regions than in other regions (e.g. annotated as “capsid proteins”, “terminases”, etc.). In order to compute each gene’s scores, input data is transformed into two sequences of indicators using data obtained from Prodigal and HMMER3 outputs. The sequence of indicators for computing “phage scores” are formed as following:

- 0 for a gene whose protein product does not match any pVOG profile HMMs
- (1 + ‘black_penalty’) for a gene whose protein product does match a pVOG profile HMM from the “black list”
- 1 for a gene whose protein product does match a pVOG profile HMM from either the “black” or “white” list

Then, a triangular window function [8] is applied to count “phage scores” using the following formula:

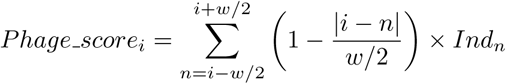

where *i* is gene index, *w* is window width, *Ind*_*n*_ - *n*-th gene’s indicator Similarly, GC scores are obtained for each gene with the following formula:

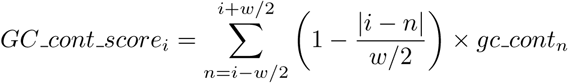

Where *gc_cont*_*n*_ is GC content for a gene obtained from Prodigal output. After the two scores are calculated, the resulting score is computed for each gene as a product of its “phage score” and “GC score”.

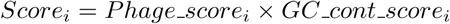

Finally, the algorithm determines phage regions based on the sequence of resulting scores. Phage regions are defined as ranges of genes with scores exceeding the “minimum score threshold”, given that at least one of the genes has a score exceeding the “maximum score threshold”. Thus, for each input contig, Phigaro returns a set of prophage regions with their coordinates. For each of the predicted prophage regions, the algorithm defines possible taxonomy of the phage and marks the possible presence of transposons inside of the prophage sequence. Phigaro produces annotated “prophage genome maps” where prophages are visualized dynamically on a webpage by displaying their proteins as arrows with colour coding of the phage functional modules (See Supplementary).

### PhigaroFinder parameters optimization

In order to optimize PhigaroFinder parameters, we used a “golden standard” set of 54 bacterial genomes with manually annotated prophage positions [3]. During a two-step optimization process, “black list” penalty, “white list” bonus, threshold values, as well as hmmscan E-value and window width were chosen. Parameter selection was done using grid search techniques and Jaccard index and PPV (Positive Predicted Value) as metrics:

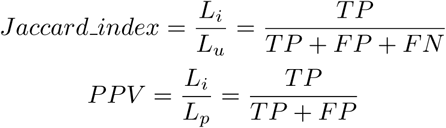

where *Li* - length of intersection of predicted and true prophage regions, *Lu* - length of union of predicted and true prophage regions, *Lp* - length of predicted phage region.

- “Black list” penalty: −2.2
- “White list” bonus: +0.7
- Minimum score threshold: 45.39
- Maximum score threshold: 46.0
- Hmmscan E-value: 0.00445
- Window size: 32 ORFs

For this set of parameters, Jaccard index was 0.625, and PPV was 0.853.

### Performance analysis

Phigaro performance was compared to those of other prophage predicting tools using prophage predictions from 54 annotated and curated bacterial genomes as input data that are commonly used for benchmarking prophage prediction tools [3]. Although there are several prophage predicting tools to date (such as Phaster [2], Virsorter [10], Phage_Finder [4], ProphET, Prophinder [7], PhiSpy [1]), only the first two accept unannotated FASTA sequences as input. To compare the performance of all of the listed tools, we used the same metrics as those used in gridsearch procedure: Jaccard index and PPV (Table 1).

**Table 1.** Performance of Phigaro compared to other commonly used prophage prediction tools

| Program | Input | App/Web | Jaccard index | PPV | Average time |
| --- | --- | --- | --- | --- | --- |
| Phigaro | .fna | standalone | 0.625 | 0.853 | 167s |
| PHASTER | .fna | web/API | 0.61 | 0.684 | 136s |
| VirSorter | .fna | standalone | 0.347 | 0.373 | 1155s |
| ProphET | .fna + .gff | standalone | 0.535 | 0.745 | 102s |
| Phage_Finder | .fna + .faa + .ptt | standalone | 0.477 | 0.626 | 238s |
| PhiSpy | .gbk/SEED | standalone | 0.377 | 0.42 | 442s |
| ProPhinder | .gbk | web | 0.341 | 0.763 | 509s |

In spite of performing less accurately than Phaster on certain bacterial genomes, Phigaro performance appears to be the best among the existing tools. The mean execution time stays in top three among all presented tools. Overall, we show that Phigaro has decent performance compared to existing prophage prediction tools. Additionally, the tool marks possible transposons inserted into prophages and provides dynamic visualizations to inspect the genome annotation and organization of prophages.

