## Supplementary for "Phigaro: high throughput prophage sequence annotation"

### Appendix: Supplementary Material

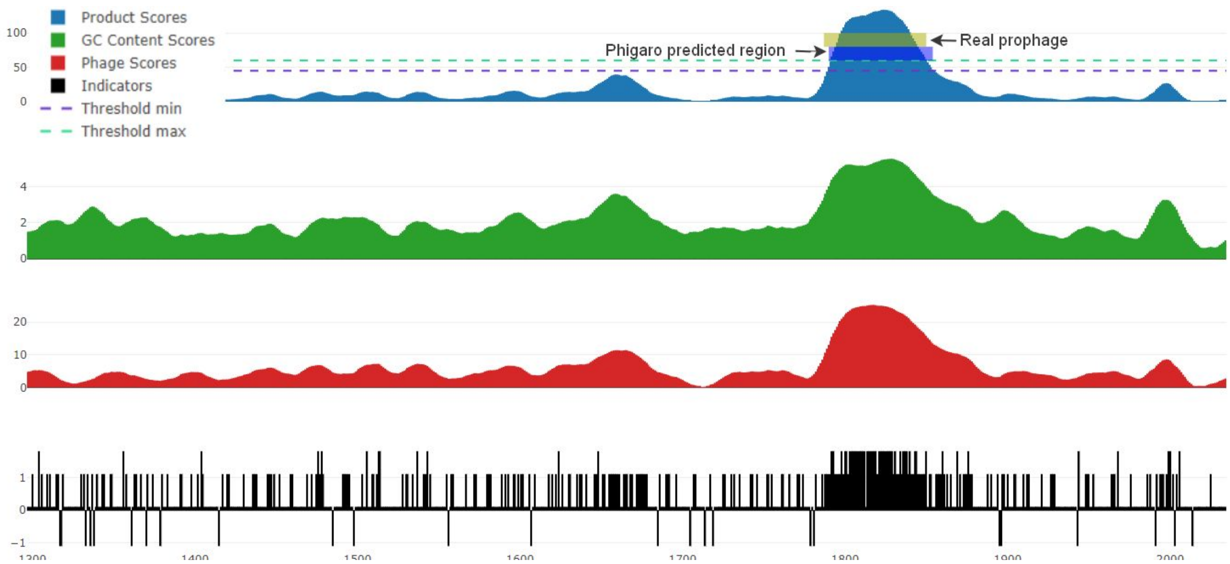


**Supplementary Figure 1**. Example of sliding window algorithm scores. Staphylococcus aureus subsp. Aureus N315 (NC_003140.1). Threshold min: 45.39, threshold max: 46.00. Top blue curve: total scores with marked predicted and real prophage regions and thresholds. Green curve: gc_content scores; red curve: phage scores. The bottom plot is a plot of indicators, based on which the phage score was calculated: the tallest black bars stand for the genes are “white phage genes” (+0.7 bonus), medium black bars are genes marked as “neutral phage genes”, the bars placed under the zero line are genes marked as “black phage genes” (-2.2 penalty) and zero bars are genes marked as “non-phage genes”.

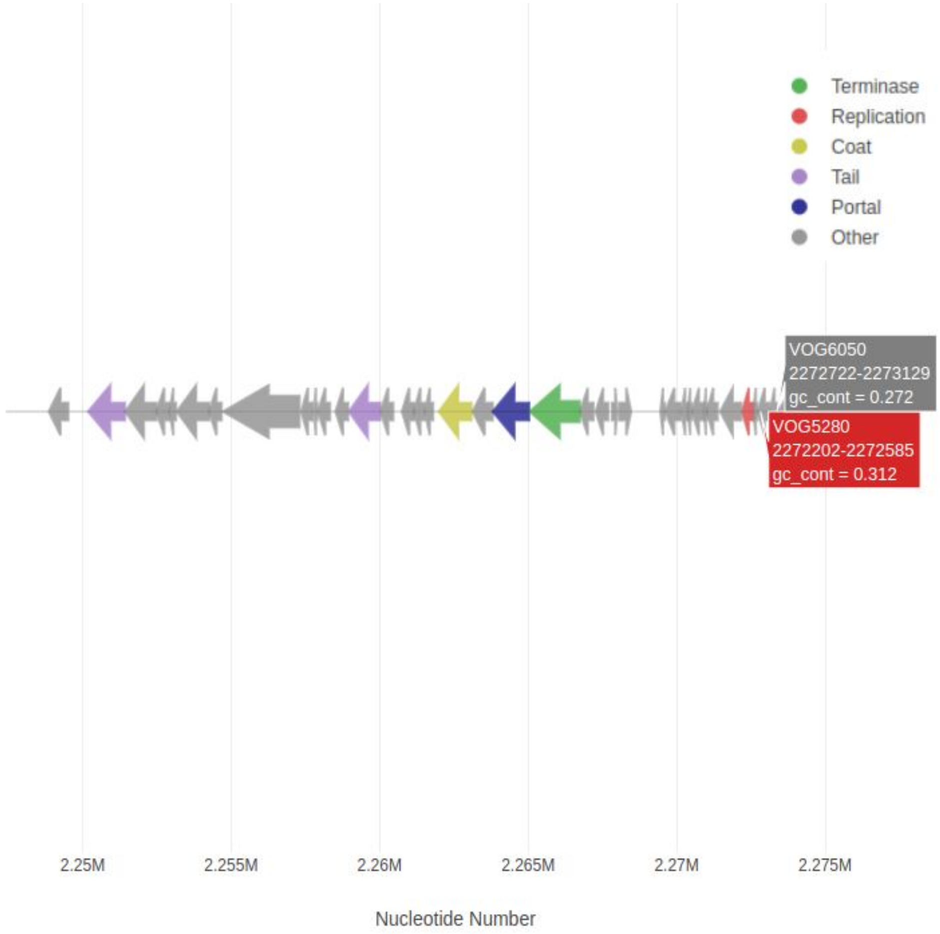


**Supplementary Figure 2.** An example of a predicted prophage genome map generated by Phigaro. Genes detected by Prodigal are presented as arrows. The direction of the arrow shows the transcription orientation. Functional categories of genes are annotated in color. The bottom axis shows the genomic coordinate (nucleotide number) along the bacterial chromosome. This is a dynamic webpage HTML interface where the user may zoom in and out of the display and click on the individual genes to view their coordinates, annotations and GC content.The user may also submit the prophage sequences directly to the online NCBI BLAST server via a single click.
